# Increasing calling accuracy, coverage, and read depth in sequence data by the use of haplotype blocks

**DOI:** 10.1101/2021.01.07.425688

**Authors:** Torsten Pook, Adnane Nemri, Eric Gerardo Gonzalez Segovia, Henner Simianer, Chris-Carolin Schoen

**Author notes:** Animal Breeding and Genetics, University of Goettingen, Albrecht-Thaer Weg 3, 37075 Goettingen, Germany (Torsten Pook).

## Abstract

High-throughput genotyping of large numbers of lines remains a key challenge in plant genetics, requiring geneticists and breeders to find a balance between data quality and the number of genotyped lines under a variety of different existing technologies when resources are limited. In this work, we are proposing a new imputation pipeline (“HBimpute”) that can be used to generate high-quality genomic data from low read-depth whole-genome-sequence data. The key idea of the pipeline is the use of haplotype blocks from the software HaploBlocker to identify locally similar lines and merge their reads locally. The effectiveness of the pipeline is showcased on a dataset of 321 doubled haploid lines of a European maize landrace, which were sequenced with 0.5X read-depth. Overall imputing error rates are cut in half compared to the state-of-the-art software BEAGLE, while the average read-depth is increased to 83X, thus enabling the calling of structural variation. The usefulness of the obtained imputed data panel is further evaluated by comparing the performance in common breeding applications to that of genomic data from a 600k array. In particular for genome-wide association studies, the sequence data is shown to be performing slightly better. Furthermore, genomic prediction based on the overlapping markers from the array and sequence is leading to a slightly higher predictive ability for the imputed sequence data, thereby indicating that the data quality obtained from low read-depth sequencing is on par or even slightly higher than high-density array data. When including all markers for the sequence data, the predictive ability is slightly reduced indicating overall lower data quality in non-array markers.

**Author summary:** High-throughput genotyping of large numbers of lines remains a key challenge in plant genetics and breeding. Cost, precision, and throughput must be balanced to achieve optimal efficiencies given available technologies and finite resources. Although genotyping arrays are still considered the gold standard in high-throughput quantitative genetics, recent advances in sequencing provide new opportunities for this. Both the quality and cost of genomic data generated based on sequencing are highly dependent on the used read depth. In this work, we are proposing a new imputation pipeline (“HBimpute”) that uses haplotype blocks to detect individuals of the same genetic origin and subsequently uses all reads of those individuals in the variant calling. Thus, the obtained virtual read depth is artificially increased, leading to higher calling accuracy, coverage, and the ability to all copy number variation based on relatively cheap low-read depth sequencing data. Thus, our approach makes sequencing a cost-competitive alternative to genotyping arrays with the additional benefit of the potential use of structural variation.

## Introduction

High-throughput genotyping of large numbers of lines remains a key challenge in plant genetics and breeding. Cost, precision, and throughput must be balanced to achieve optimal efficiencies given available technologies and finite resources. Improvements of the cost-effectiveness or resolution of high-throughput genotyping are a worthwhile goal to support efforts from breeders to increase genetic gain and thereby aiding in feeding a planet with a rising Human population [1].

Currently, high-throughput genotyping is performed using single nucleotide polymorphism (SNP) arrays in most common crops and livestock species. These arrays can have various densities, ranging from 10k SNPs [2] to 50k [3] to 600k SNPs [4,5], are relatively straightforward to use [6], and typically produce robust genotypes with relatively few missing calls or calling errors [5]. As a result, genotyping arrays are widely used for a broad range of applications, including diversity analysis [7,8], genomic selection [9,10] or genome-wide association studies [11,12]. Limitations of the technology comprise the complexity and cost of designing the arrays, their inability of typing *de novo* polymorphisms, their lack of flexibility in the choice of marker positions, and the cost of genotyping which increases significantly with the number of SNPs on the array. In addition, array markers are typically SNPs selected to be in relatively conserved regions of the genome [13,14], i.e. by design they provide little information on structural variants, although calling of structural variation is also possible via genotyping arrays [15].

In recent years, rapid advances in next-generation sequencing (NGS) have enabled targeted genotyping-by-sequencing (GBS) and whole-genome-sequencing (WGS) to become cheaper, more accurate, and widely available [16,17]. Compared to genotyping arrays, GBS and WGS data provide additional information like the local read-depth and a higher overall marker density that have been successfully used in a variety of studies [18–20]. Studies that use GBS or WGS data to call structural variation typically use a read-depth of at least 5X [21]. For applications such as genomic prediction, the use of 1X to 2X read-depth would be imaginable. However, as of today, reported prediction accuracies when using plain sequence data in such approaches are substantially lower [22]. With known pedigrees [23] and/or founder lines with higher read-depth [24] even a lower average read-depth was shown to be useful for genomic prediction, although the predictive ability is still slightly below that of array data. A key limitation of NGS is, that the cost of sequencing increase almost linearly with the sequencing depth [25]. Thus, generating sequence data with adequate read-depth is still too costly for most routine applications and thus genotyping arrays are still considered the gold standard in high-throughput quantitative genetics.

Importantly, due to stochastic aspects of sequencing in sampling from genomic reads, not all variants are called in whole-genome sequencing at low to very-low depth (e.g. below 1-2x) [6, 26]. In the context of a sequenced population, virtually every variant position displays significant amounts of missing calls, leaving these gaps to be filled prior to subsequent applications. This *in silico* procedure is referred to as imputation. Over the years a variety of approaches for imputation have been proposed [27–31]. The interested reader is referred to Das et al. [32] for a detailed review and comparisons between commonly used imputation software. As tools are typically developed for application in human genetics with high genetic diversity, parameter optimization is mandatory for livestock and crop populations [33]. However, as long as somewhat related individuals are considered and parameter settings are chosen adequately [33], error rates for imputation of array data are usually negligible.

One of the limitations of imputation when working with low read-depth sequence data has been the challenge of phasing reads, causing imputation error rates to increase notably. In contrast to human and livestock genetics, where phasing is an absolute requirement, fully inbred and homozygous lines are readily produced in maize [7,34] and other plant species [35]. Such inbred lines are becoming more used in breeding to, among others, reduce the length of the breeding cycle, increase the genetic variance and safeguard genetic diversity [7,36–38]. Without the need for phasing, there is high potential in using very-low to low sequencing depth to genotype a large number of lines and apply efficient imputation to obtain maximum data quality at minimal cost. Specifically, information on read-depth could be used to support imputed variant calls. To our knowledge, none of the existing imputation approaches currently addresses this.

In this work, we propose a new imputation pipeline (“HBimpute”) for sequence-derived genotypic data of homozygous lines that uses long-range haplotype blocks from the software HaploBlocker [39]. Haplotype blocks in HaploBlocker indicate cases of group-wise Identity-by-descent (IBD) [40]. This information serves to artificially merge reads of lines in the same haplotype block to locally increasing the read-depth allowing higher calling accuracy and thereby increase calling precision and reduce the share of missing calls. We compare our imputed data to array data, high read-depth sequence data, and low read-depth sequence data that were imputed via the state-of-the-art software BEAGLE [31]. The performance of the different datasets is evaluated by their respective usefulness in a subsequent genome-wide association study (GWAS) and for genomic prediction (GP).

## Results

In the following, we will briefly sketch the key steps of the HBimpute pipeline (Figure 1). As a first step of the pipeline, read-mapping and variant calling are performed to generate a raw SNP-dataset with a potentially high share of missing calls. For this, we suggest the use of FreeBayes [41], but software such as GATK [42] and a workflow along the GATK best practices [26] is a valid alternative here.

**Fig 1.**
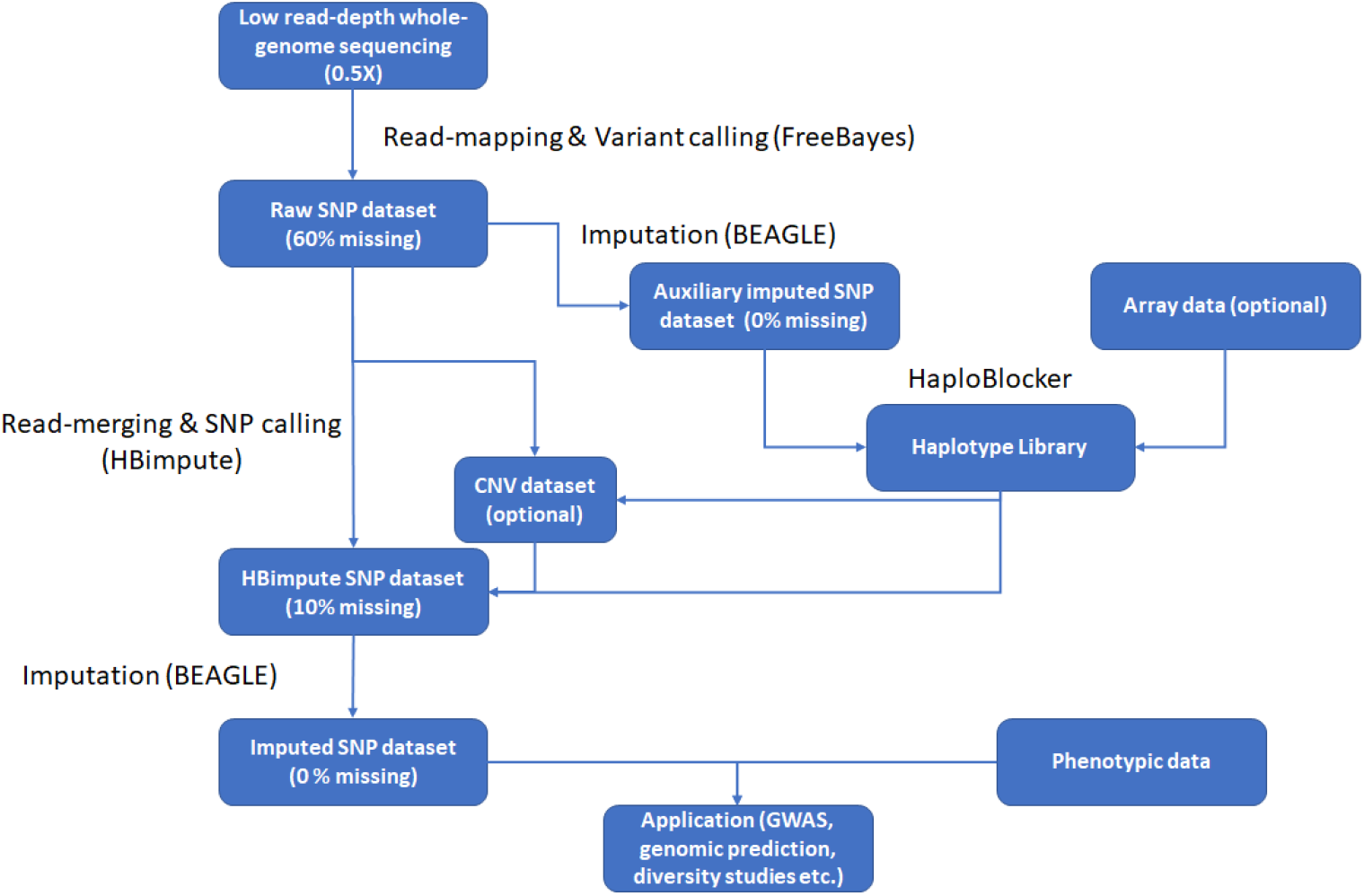
Schematic overview of the HBimpute pipeline with exemplary data attributed (read-depth / share missing) according to the maize data.

Secondly, a haplotype library for the dataset at hand needs to be derived via the software HaploBlocker [39]. As HaploBlocker is not supporting a high share of missing data, one first has to generate an imputed dataset (auxiliary imputed SNP dataset, Figure 1) and use this set for the calculation of the haplotype library. A potential software to use here is BEAGLE [31]. Instead of using the sequence data itself, the haplotype library can also be computed from other genotypic data of the considered lines like array data. In the following, we will present results for two approaches and distinguish between HB-seq and HB-array, depending on whether the sequence data itself or 600k array data [5] was used to derive the haplotype library.

Thirdly, the information regarding local IBD from the resulting haplotype library is used in a second variant calling step. In contrast to the initial variant calling, all mapped reads from individuals that are locally in the same haplotype block are also used for each respective individual. As the local read-depth in most regions is massively increased via the local merging procedure an optional step to detect copy-number-variation (CNV) can be performed. Lastly, the resulting dataset (HBimpute SNP dataset, Figure 1) is imputed via traditional imputing software (imputed SNP dataset, Figure 1) [31] and can be used for subsequent downstream applications.

We applied our imputation pipeline on a dataset of 321 maize doubled haploid lines (DH), derived from an open-pollinated landrace [43]. The DHs were whole-genome sequenced at 0.5X read-depth with 2,152,026 SNPs being called by FreeBayes [41] (compared to 616,201 on the high density array [5]). Even though the differences in marker density between the sequence and array data are going down slightly after applying quality control filters, removal of fixed markers, and imputation (1,069,959 vs 404,449 SNPs), this still is a substantial increase in marker density.

When using the HB-seq pipeline, the average read-depth increased from 0.53X to 83X. As a result, the share of cells of the genotype dataset that were called increases from 39.3% before merging to 95.2% after haplotype-block merging. Note however that the read-depth varied greatly between lines and genomic regions, as it depends primarily on the frequencies of haplotype blocks in the population. When using HB-array an average read-depth of 51.3X was obtained with 93.1% of the variants being called. This smaller increase in average read-depth is mostly due to longer haplotype blocks fewer individuals being identified in HaploBlocker. However, lower read-depth not necessarily means lower data quality in this instance, as higher relatedness between lines in the same haplotype block can reduce noise introduced by individuals that are similar but not the same locally. In fact, we expect the quality of the array-based haplotype library to be higher than the one obtained via BEAGLE imputed low read-depth sequence data (HB-array) as the share of missing calls in the raw array data is substantially lower (1.2% vs. 60.7%) [33]. In practice, such data is however usually not available when sequence data is generated. Parameter settings in HaploBlocker can be adjusted to control the structure of the haplotype library [39].

## Imputation

When comparing discordance rates of the imputed SNP dataset with the genotype data from the 600k Affymetrix® Axiom Maize Genotyping Array [5], error rates overall are reduced from 1.03% to 0.60% in the HB-seq pipeline and 0.50% in the HB-array pipeline (Table 1). Error rates here refer to discordance rates between the respective imputed panel and the 600k data. The dataset was split into three classes to further access the performance of the imputation (Figure 1):

1. Cells first called in FreeBayes step (“Present in raw-data”)
2. Cells first called in HBimpute step (“With call after HB”)
3. Cells first called in the imputed SNP dataset (“Without call after HB”)

**Table 1.**
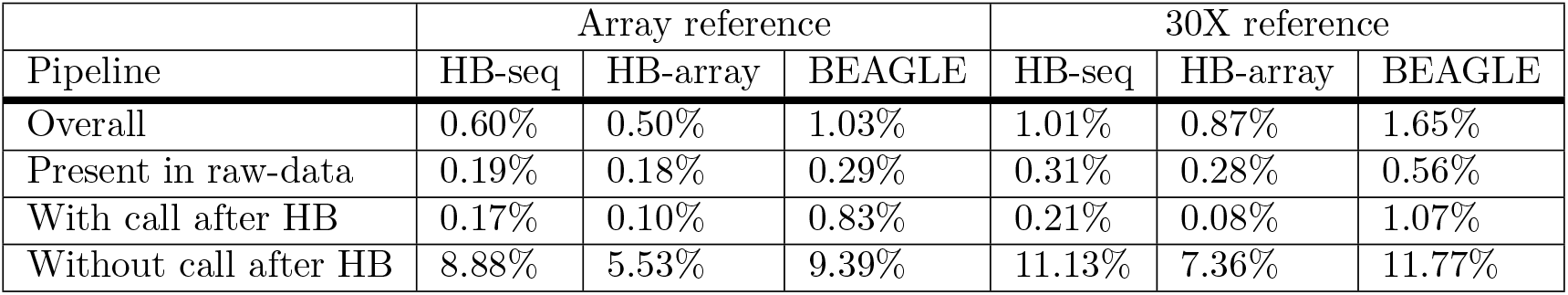
Discordance rates of the imputed sequence data and the comparison data set (array / high read-depth sequence data) for different imputation algorithms.

| Pipeline | Array reference |  |  | 30X reference |  |  |
| --- | --- | --- | --- | --- | --- | --- |
|  | HB-seq | HB-array | BEAGLE | HB-seq | HB-array | BEAGLE |
| Overall | 0.60% | 0.50% | 1.03% | 1.01% | 0.87% | 1.65% |
| Present in raw-data | 0.19% | 0.18% | 0.29% | 0.31% | 0.28% | 0.56% |
| With call after HB | 0.17% | 0.10% | 0.83% | 0.21% | 0.08% | 1.07% |
| Without call after HB | 8.88% | 5.53% | 9.39% | 11.13% | 7.36% | 11.77% |

For all three classes improvements in calling accuracy are obtained with the highest gains for those cells that were first called in the HBimpute step, as the average error rate is reduced from 0.83% to 0.17% / 0.10% in HB-seq / HB-array. Similarly, discordance rates for cells already called in the FreeBayes step are reduced by about 40% as calls are overwritten (0.29% vs. 0.19 / 0.18%, Table 1) when a high number of other individuals in the same block carry the other variant, thus indicating the power of our approach to detect calling errors. Note that the imputed dataset in HB-array was compared to the same array data that was used for the calculation of the haplotype library and therefore results for HB-array are potentially downward biased. However, as similar improvements were observed when comparing the imputed data panel to high read-depth sequence data this effect should be negligible. Due to the overall higher data quality and lower share of missing markers after the HBimpute step, even error rates for cells imputed in the subsequent BEAGLE imputation step are reduced slightly.

When comparing discordance rates of the imputed sequence data to the 30X sequence data that was generated for seven of the lines considered, we again observe much better results in the dataset imputed via our suggested pipeline (HB-seq: 1.01% / HB-array: 0.87%) compared to the dataset imputed via BEAGLE (1.65%, Table 1). In contrast to the comparison with the array data, error rates for positions called in the HBimpute step are even lower than for markers called in the FreeBayes step, as overwriting of already called variants requires stronger evidence than calling a previously missing variant. Even though overall error rates seem to be higher when comparing to the high read-depth sequence data, this is mostly due to lower overall error rates in SNPs that were placed on the array. When just considering marker positions that are also on the array error rates reduce to 0.90% for HB-seq, 0.68% for HB-array, and 1.38% for plain BEAGLE imputation [31]. Cells with no called variant in the 30X sequence data were ignored here.

When analyzing the allele frequency spectrum of the resulting data panel (Figure 2.A-D), we can observe an increased number of markers for all minor allele frequencies. Overall, the distribution of the allele frequency spectrum looks very similar with an approximately three-fold increase in the number of markers for the sequence data. When just considering marker positions that are overlapping with the 600k array, a higher share of extremely rare variants (¡1%) in the sequence data can be observed (Figure 2.E-H). As the minor variant is more difficult to impute and for the 600k data 98.8% of all variants were called before imputation [33] this distortion to the more frequent variant should be expected. The total number of non-fixated markers that are shared between array and sequence is lowest in the 600k array data with only 366,822 SNPs compared to 368,095 SNPs for HB-seq, 369,211 SNPs for HB-array, and 377,900 SNPs for plain BEAGLE imputation. The data panel created by imputation via BEAGLE is including the highest number of rare variants, which is also due to rare variant calls being overwritten in HB-seq and HB-array when strong support for the alternative is given.

**Fig 2.**
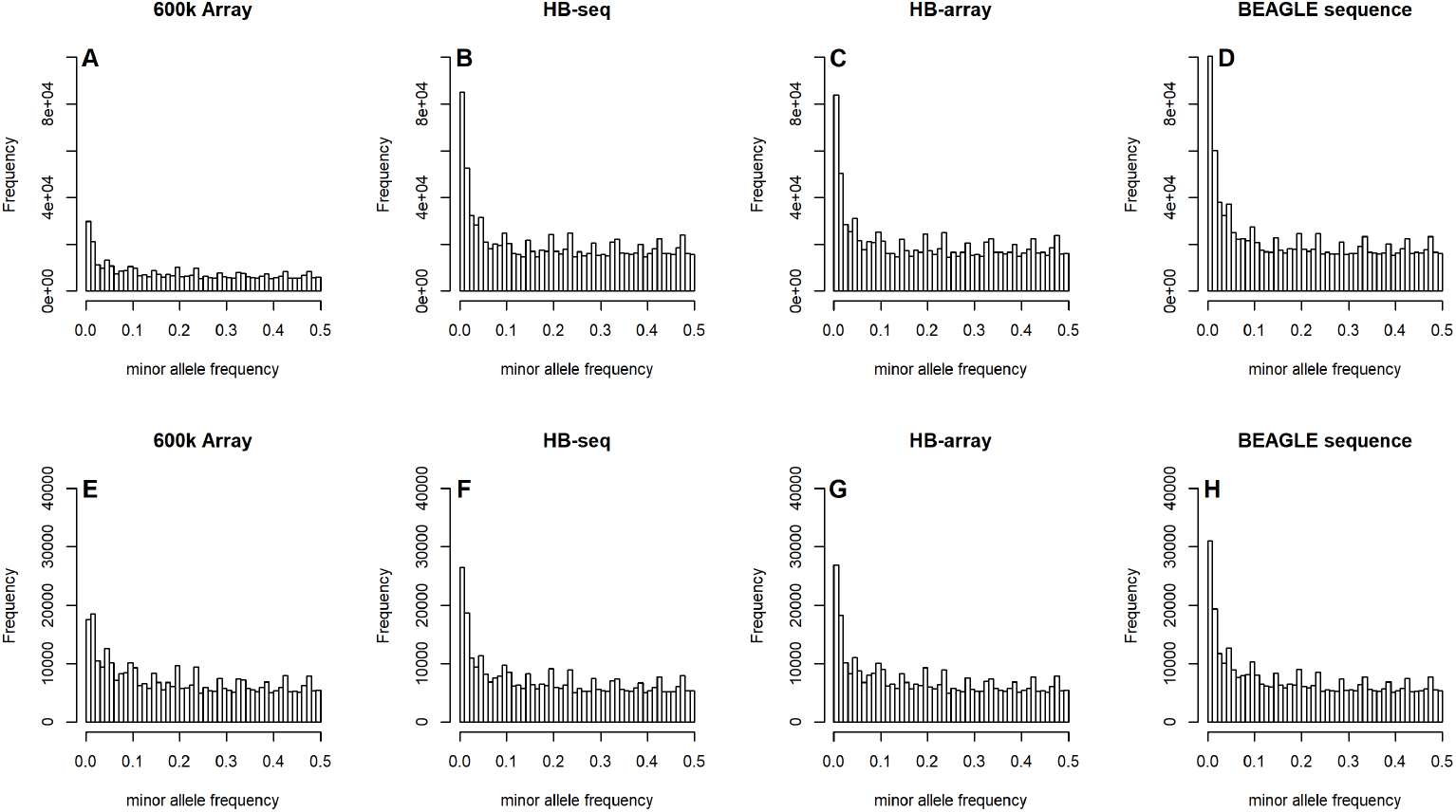
Allele frequency spectrum of the different genotypic datasets for all markers (A-D) and the panel of markers shared between array and sequence data (E-H).

### Estimation of local read-depth and structural variation

Genotyping of structural variants from read-mapping typically requires higher sequencing depth than calling SNPs. When comparing the obtained locally smoothed read-depth of the 30X sequence data to the imputed low sequence data, we observed an average correlation of 0.750 compared to 0.257 for the raw 0.5X data, indicating that the imputed data can be used for the calling of structural variation (correlation without local smoothing: 0.442 vs 0.102). Visual inspection of local read-depth also shows that peaks (Figure 3.A/C) and local pattern (Figure 3.B/D) between the imputed low read-depth sequence data and the high read-depth sequence data match, whereas the raw low read-depth sequence data has much higher volatility (Figure 3.E/F). Note that HBimpute can only provide an estimated read-depth for regions that are in a local haplotype block, leading to some gaps (4.1%, Figure 3.C/D).

**Fig 3.**
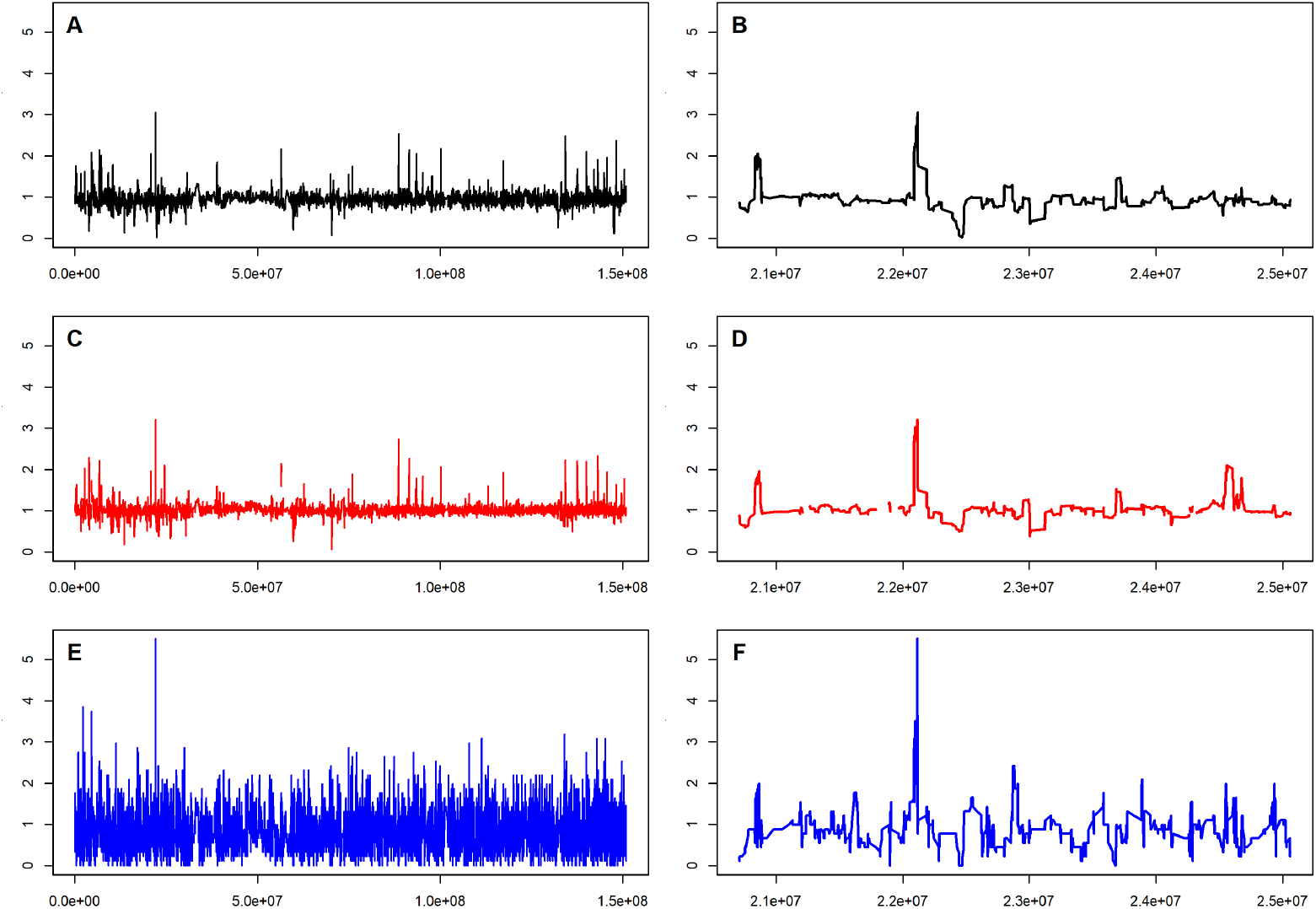
Estimated standardized read-depth for line PE0213 via the use of high read depth sequence data (A/B), imputed low read-depth sequence data via HBimpute (C/D) and raw low depth depth sequence data (E/F) for chromosome 10 and an exemplary chosen segmented in a peak region.

### Genomic prediction

The performance of the HBimpute and BEAGLE imputed sequence data for genomic prediction was evaluated against that of the array data. For this, we compared the obtained predictive ability of each set for nine traits, including early vigor and plant height at different growing stages, days to silking, days to tassel and root lodging [43]. The predictive ability for the imputed sequence data panels was marginally lower for eight of the nine considered traits. Differences between data panels were however small as the average difference was only 0.0028 and at most 0.0069 (Table 2 & Supplementary Table S1). On the contrary, when using only the markers positions that are shared between the sequence and the array data, minor improvements were obtained for eight of the nine traits (paired t-test, p-values ¡ 10^−15^). As differences on average are just 0.0011 this should still be neglectable in practice. Including CNV calls from the HBimpute pipeline led to slightly reduced predictive abilities, in particular when applying no filtering.

**Table 2.** Average predictive ability for the nine maize traits [43] depending on the genotype data used for prediction. The panel of overlapping markers includes all markers included in the array and sequence data panel after quality control filtering.

| Pipeline | HB-seq | HB-array | BEAGLE | HB-seq + CNV | HB-array + CNV | Array data |
| --- | --- | --- | --- | --- | --- | --- |
| Predictive ability | 0.5152 | 0.5156 | 0.5151 | 0.5096 | 0.5114 | 0.5180 |
| Predictive ability (overlap) | 0.5191 | 0.5194 | 0.5186 | 0.5170 | 0.5187 | 0.5183 |

### Genome-wide association study

Furthermore, we evaluated the suitability of the imputed low read-depth sequence data to be used in a GWAS, compared to high-density array data. Our goal was to estimate whether the higher number of variants genotyped compared to the chip impacted the power or resolution of GWAS. When comparing the Manhattan plots derived based on sequence data and array data on simulated traits, in general, higher peaks are observed for all variants of the sequence data, leading to a higher number of regions identified when using the same p-values. As GWAS results for different data sets and the same p-values are not comparable and correction of a significant threshold is not straightforward, we instead report the share of true positive QTL hits compared to the total number of regions with a GWAS hit.

Best results were obtained with sequence data imputed via HB-array and plain BEAGLE imputation of the low read-depth sequence data, closely followed by the 600k array data and HB-seq (Figure 4.A, Supplementary Table S2). Filtering of sequence data, as done for genomic prediction, leads to increased false discovery rates. Even higher drops in performances were observed when using a genotyping array with 10k / 50k markers. Overall, results for the different imputed low read-depth sequence data panels are very similar and data panels with a high number of markers seem most suitable for a GWAS analysis. However, applying less strict quality control filters to increase the number of included SNPs to 1,349,597 SNPs did not improve GWAS results for HB-seq, thus showing that data quality still should not be neglected.

**Fig 4.**
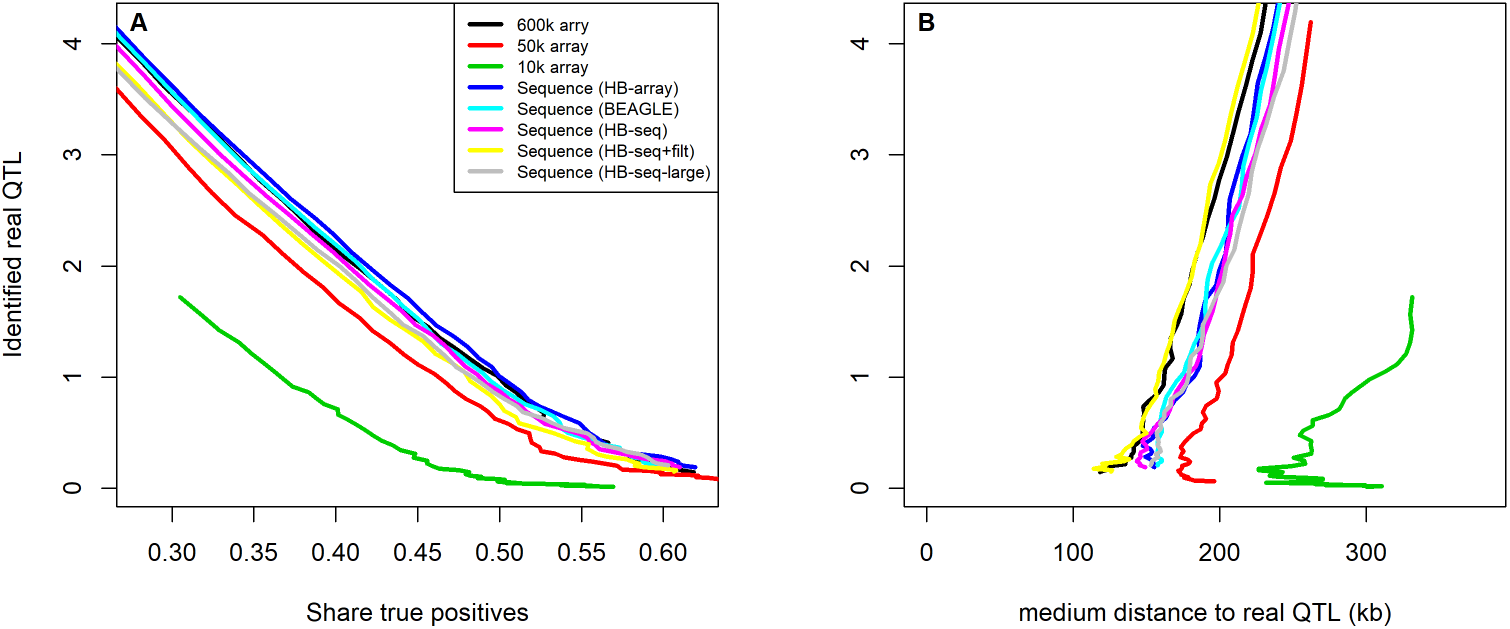
Number of positive GWAS hits for simulated traits with 10 underlying QTL according to the share of true positives (A). Median distance of the local GWAS peak (highest p-value) and the underlying true QTL for true GWAS hits (B).

In terms of mapping power, we observed the lowest median distance between the GWAS peak (highest local p-value) and the underlying true QTL when using the 600k data and the HB-seq data when only including markers shared with the array (Figure 4.B). This indicates that for fine-mapping the quality of markers should be more important than the total number of markers. Nonetheless, down-sampled array data (10k / 50k) again performed worst. For all sequence-based data panels, a relatively high number of isolated GWAS hits were observed that could potentially be due to transposable elements and other structural variation as the B73 reference genome [44, 45] represents dent germplasm whereas the lines in this study belong to the flint pool [46].

## Discussion

HBimpute is shown to be a pipeline for accurate imputation of low read-depth sequence data. Results indicate that the use of HBimpute allows sequencing at reduced depth while maintaining data quality comparable to high-density array data. Thus, HBimpute leverages significant cost savings and/or higher quality data for subsequent applications.

Overall, we can confirm that WGS and GBS are valid alternatives to genotyping arrays for the generation of genotypic data and use in subsequent applications. For GWAS the use of genomic data generated via sequencing has been shown to be more powerful than the use of traditional array data, even though differences between the sequence data and the 600k data were relatively small. Although error rates of the imputation were reduced by HBimpute, differences in the performance of the GWAS analysis were neglectable compared to when using a BEAGLE imputed dataset, and the improved performance can mostly be attributed to an increased marker density [12]. Even though the predictive ability on the full data panel for the sequence data was slightly lower than when using array data, this can mostly be attributed to the inclusion of some low-quality markers in the sequence data. This effect is even stronger when including CNV calls. When just considering genotype information on the panel of overlapping markers between sequence and array data the predictive ability was marginally improved, indicating that the overall data quality of low read-depth sequence data is on par or even slightly higher than array data. This is further supported by higher imputing error rates on non-array markers and slightly increased predictive ability when using HBimpute instead of BEAGLE for sequence imputation. Although no improvements for genomic prediction or GWAS were obtained by accounting for structural variation or including non-array markers, there should still be at least some high-quality markers and functional CNVs not captured by the array. In particular when analyzing specific regions of the genome or by applying improved quality control filtering, this information can still be valuable.

Overall, we can conclude that rating the usefulness of a genomic dataset is dependent on the application at hand and data preparation and filtering should be chosen accordingly. With rising marker density calling and imputing error will increase (due to the inclusion of low-quality markers) and an adequate weighting between marker density and quality has to be found. E.g. when conducting a GWAS focus should be on including a high number of markers, whereas for genomic prediction high-quality markers have shown to be more important in this study. In this context HBimpute is providing a framework to improve imputation accuracy and thereby improve data quality compared to existing imputation software. Note that both GWAS and genomic prediction via a mixed model are quite robust methods that will neutralize most of the issues of partially poor data quality.

The use of sequence data comes with both challenges and opportunities. Sequence data is providing more information in less conserved regions and thus is providing more information on structural variation of the genome [47]. In particular crop genomes tend to have a high share of transposable elements (e.g. 85% in maize [44]). As data in those regions will always be noisier than array markers that are specifically selected to be in more conserved regions [5,13]. Note that high-quality genotyping arrays are not available for all species and the relative cost of sequencing will be lower for species with short genomes.

Therefore, the decision on which genotyping technology to use in practice will be highly dependent on the species at hand, its genome length, available genotyping arrays, and intended subsequent applications. Even though it is possible to replace BEAGLE with other imputing software, other tested tools either needed more computing time, were not as accurate for DH-lines, or struggled with a high share of missing data, leading us to highly recommend the use of BEAGLE [31].

A key limitation of the HBimpute pipeline is that it requires highly accurate phase information that is typically not obtainable for low read-depth sequence data in non-inbred material and therefore is mainly applicable to inbred lines. However, with the availability of long-read sequencing technologies and highly related individuals with available pedigree information, as commonly present in livestock genetics, this might change in the future. The here proposed HBimpute pipeline and software can be applied on heterozygous data in the same way as with inbreeds by handling the two haplotypes of each individual separately.

In particular for the detection of structural variation, the here suggested pipeline is shown to be highly efficient, as local read-depth of the imputed 0.5X data was very similar to 30X data that was generated for seven of the studied lines. Through this, we conclude that the use of HBimpute can enable the calling of structural variation from 0.5X data. As other studies detecting structural variation typically rely on 5X or even 10X data, this is a massive cost reduction, thereby enabling the calling of structural variation on large-scale populations.

## Materials and Methods

In the following, we will describe the haplotype block-based imputation step of our proposed pipeline in more detail. This step is typically applied after an initial SNP calling step that is resulting in a dataset, we refer to as the raw SNP dataset (Figure 1). In our test each of the 340 individual DH lines had its raw read file (FASTQ) aligned to the B73v4 reference genome [48] using BWA MEM [45]. Subsequently, variant calling in FreeBayes was performed using 100 kilo-base pair genome chunks with marker positions from the 600k Affymetrix® Axiom Maize Genotyping Array [5] given as input to force variant reporting at those locations (−). Furthermore, 5 supporting observations were required to be considered as a variant (−C 5) with at most 3 alleles per position (−use-best-n-alleles 3) and a maximum total depth in a position of 340 (−max-coverage 340). To ensure adequate data quality, markers with more than 1% heterozygous calls are removed since we would not expect heterozygous genotypes for DH lines.

Subsequently, 19 lines were removed from the panel, as genotypes from the 600k array and sequence data showed strong indication for contamination and/or mislabeling (see Genotype data used subsection).

The newly proposed HBimpute step is using the raw SNP dataset (Figure 1) as the only mandatory input and can be separated into three sub-steps, that will be discussed in the following subsections:

1. Derivation of a haplotype library
2. Read-merging
3. SNP-calling

Note, that only those reads used for the variant calling in the Variant call format (VCF) - file are used and in particular that in no step of the proposed algorithms original raw read data from the Binary Alignment Map (BAM) - files or similar need to be accessed. After executing these steps, the resulting HBimpute SNP dataset (Figure 1) is obtained, with only a few remaining missing calls. Nonetheless, subsequent imputation via traditional imputation software is necessary for most applications. In our tests, the software BEAGLE performed well both in terms of computing time and accuracy [31] and was chosen for all reported tests. We will here focus on describing the default settings of the associated R-package HBimpute, but also discuss potential deviations with most parameters in the tool being adaptable to set a weighting between imputation quality, number of markers considered, and the overall share of markers called in HBimpute.

Individual steps of the procedure will be explained along the example dataset is given in Figure 5 with five haplotypes and ten markers each. For simplicity, we are here assuming a read-depth of one for all called genotype entries.

**Fig 5.**
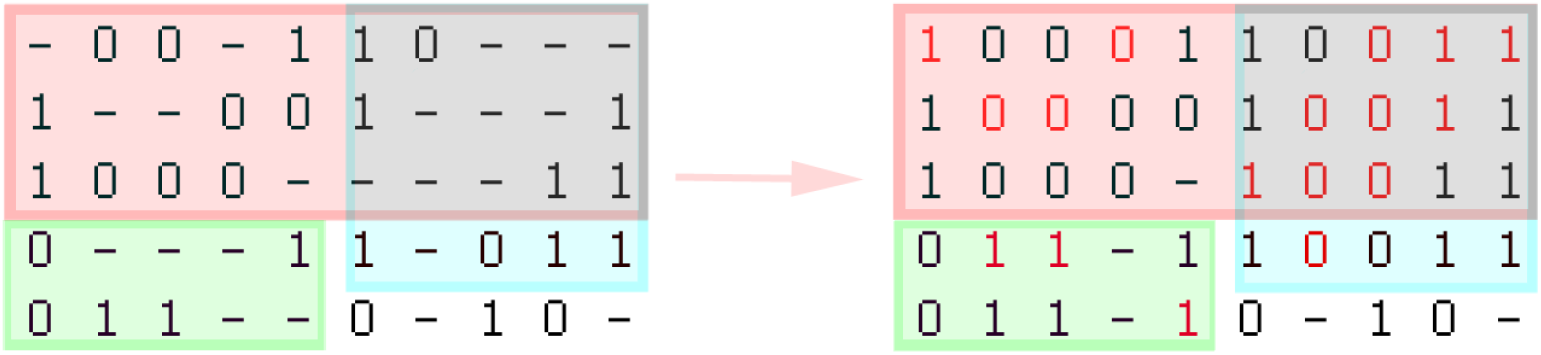
Toy example for the HBimpute step. Each column is representing a SNP and each row is representing a haplotype (for inbreds: individual). Haplotype blocks are indicated by colored blocks. Note that the blue and red block are overlapping.

### Derivation of the haplotype library

In the first step of the HBimpute, the objective is to derive a haplotype library via the associated software HaploBlocker [39]. As HaploBlocker itself is not supporting a high share of missing data, the raw SNP dataset first needs to be imputed to generate an auxiliary imputed SNP dataset (Figure 1). Alternatively, other genetic data of the considered lines like array data can also be used. Results for both approaches (HB-seq & HB-array) are presented in the results section. Since the overall data quality in our tests indicates that the array data is slightly higher quality in terms of consistency and overall calling error rates than the raw low read-depth sequence data, the use of array data is recommended when available (HB-array). Furthermore, additional lines can be included as a reference panel in both approaches. Individuals in the reference panel can either be used to improve the quality of the haplotype library and/or provide additional reads to be used in the subsequent read-merging step. In all our tests, the default settings in HaploBlocker were adjusted to identify long haplotype blocks which are potentially present in low frequency (node_min = 3, edge_min = 3, weighting-length = 2 [39]) and a target coverage was set to ensure sufficient coverage of the haplotype library (target-coverage = 0.95 [39]). For datasets with more variations a reduction of the window size might be needed to detect shorter haplotype blocks, which can be obtained by activating adaptive window sizes (adaptive_mode = TRUE [39]), but comes with a substantially increased computing time.

For our toy example given in Figure 5 three blocks are identified with the red block including haplotypes 1,2,3 spanning over SNPs 1-10, the green block including haplotypes 4,5 spanning over SNPs 1-5, and the blue block including haplotypes 1,2,3,4 spanning over SNPs 6-10.

### Read-merging

The output of HaploBlocker is a haplotype library. As contained haplotype blocks are representing cases of group-wise IBD [40] this means that all included haplotypes should have locally matching sequences and that all reads of these lines can be used for the subsequent SNP-calling. To still be able to detect recent and rare variation, the reads of the individual itself are used with a higher weighting in subsequent steps (default: five times as high). Variant calls that were missing in the initial variant calling in

FreeBayes [41] and were only imputed in the step of the derivation of the haplotype library are ignored in this step. In our example, this means that for marker 1 in haplotype 1 there are no reads supporting variant 0 and two reads supporting variant 1. Similarly, for marker 5 there are five reads supporting variant 1 and only one read supporting variant 0 as the read of the haplotype itself is counted with a higher weighting. Note that in a real haplotype library each block usually contains far more haplotypes with the default minimum being 3 and therefore a much lower relative weighting is put on the haplotype itself.

### SNP-calling

After the read-merging step, a further SNP calling step is necessary. Since it is neither possible nor necessary to obtain calls for all markers in this step, the focus is on retrieving calls for markers with clear evidence of a certain variant. In our case, this means that at least 80% of all reads are supporting the same variant. In case no call was obtained in this step, but a variant was called in the original raw SNP dataset, this variant is inserted. This is mainly done to avoid losing rare variants.

In the toy example (Figure 5), in marker 5 variant 1 is called for haplotype 1 as five of the six reads considered support variant 1. Even though haplotype 2 is in the same local haplotype block variant 0 is called here, as the reads of the line itself are weighted higher. For haplotype 3 no variant can be called as both variants are supported by exactly one read, thus not exceeding the 80% threshold.

### Quality filters

To ensure per marker data quality, all markers with an estimated read-depth that is below 50% of the overall mean read-depth are removed from the dataset. Similarly, all markers with more than 50% missing calls are removed. These settings can be seen as relatively conservative as only markers with extremely low call rates are removed. Thus the introduction of potential noise from low-quality markers in the subsequent BEAGLE imputation procedure is reduced. Further increasing filter thresholds will increase calling precision but also potentially lose usable information.

### Optional: CNV-calling

As the read-depth after the HBimpute-based SNP-merging is massively increased the SNP-calling step can be combined with an optional step to detect structural variation, and in particular CNVs. To negate issues of high per marker variance in read-depth, we first apply a kernel smoothing function to estimate the local read-depth of the population. This is done via a Nadaraya-Watson-estimator [49] with a Gaussian kernel and set bandwidth (default: 0.25 Mega-base pairs (MBp). The local read-depth of a single haplotype is then compared to the population average with regions above 1.3 of the expectation being classified as CNVs and regions below 0.7 being classified as deletions. By adjusting the bandwidth of the smoothing function the resolution of the identification can be adapted to specifically target short/long CNV segments. Note that this approach will not detect other structural variation such as translocations, inversions, or insertions as not all raw reads from the BAM file, but only aligned reads that were used for the variant calling in the VCF-file are used here. Note that instead of performing the HBimpute step on the VCF-file, merging could also be directly applied to the reads themselves, followed by a second run of a variant caller.

For simplicity reasons in the toy example (Figure 5), we are assuming here that only the marker itself is impacting the CNV method proposed and thus no local smoothing is applied. This would result in the average read-depth in marker 4 being 0.4X (two read for five haplotypes). Haplotypes 4,5 have an estimated read-depth of 0 as no variant was called. Haplotype 1 has an estimated read-depth of 0.285X (two read for seven haplotypes) as the haplotype itself is counted five times. Both Haplotype 2 and 3 have an estimated read-depth of 0.857X (six reads for seven haplotypes). This would lead to deletions being called for haplotypes 4 and 5 (0X / 0.4X ¡ 0.7) and duplications being called for haplotypes 2 and 3 (0.857X / 0.4X ¿ 1.3). Note that this small-scale toy example is not constructed for the identification of CNVs and a much higher number of supporting reads and local smoothing is usually required for the detection of copy number variation. Both deletions and duplications are thereafter added as an additional binary marker that is coding if the respective structural variation is present in each marker or not.

Other single SNP or window-based approaches on the read-depth were also tested [50], but had limited success. No testing has been done with split read or assembly approaches [51] as all analysis in HBimpute is just using the VCF-file as input. Methods should however be relatively easily extendable to such approaches to enable the detection of other types of structural variation.

### Heterozygous data

In principle, the same pipeline suggested for inbreds can also be applied on diploid and heterozygous data that is using the two respective haplotypes separately. However, as the phasing accuracy of low read-depth sequence data is usually relatively low, the derivation of an accuracy haplotype library is heavily impacted by the software used for the initial phasing, leading to results of the SNP-calling to be very similar to the original phased and imputed datasets from the respective external software. With advanced in long-read sequencing the phasing quality might improve in the future.

### Genomic prediction

The usability of the different datasets for genomic prediction was evaluated by comparing each set for its predictive ability for nice real phenotypes, including early vigor and plant height at different growing stages, days to silking, days to tassel, and root lodging. For this, the dataset was split into 280 lines used for training and 41 lines as the test sets, and the correlation between underlying phenotypes of the test set and their estimated breeding values was used for the evaluation. We define the predictive ability as the correlation between the estimated breeding values and the phenotypes in the test set. For the evaluation a linear mixed model [52] with genomic relationship matrix [53] was used (genomic best linear unbiased prediction). This procedure was repeated 250 times for all considered traits.

### Genome-wide association study

To compared the performance of the different obtained imputed datasets a genome-wide-association-study on simulated phenotypes and therefore known underlying regions was conducted. For each trait 10 underlying QTL were simulated with 5 QTL positions randomly drawn and evaluated based on the 600k data and 5 QTL positions drawn and evaluated based on the HB-seq data. Heritability *h*^2^ of the simulated traits was assumed to be 0.5 with all 10 QTLs having equal effect size. In addition to the imputed datasets, the 600k array data was also downsampled to artificially generate low-density SNP arrays (10, 50k). Markers above a certain p-value were put in a joined region in case they are at most 1 MBp apart from each other and a region was considered a positive hit in case the underlying QTL was at most 1 MBp away from the region. The given procedure was repeated for 5,000 separately simulated traits.

### Genotype data used

For all tests performed in this study low read-depth sequencing data with a target read-depth 0.5X was generated for 340 maize doubled haploid lines, derived from an open-pollinated landrace [43]. Variants were called used the software FreeBayes [41] with marker positions of the 600k Affymetrix® Axiom Maize Genotyping Array [5] being forced to be called. This resulted in a data panel of 2,152,026 SNPs and an average read-depth of 0.73X. 19 lines were removed from the panel as genotype calls between the called variants and independently generated data from the 600k array [43] differed by more than 0.75% indicating sample contamination. Furthermore, re-labeling of 4 lines was performed as genotypes were matching with different lines based on the 600k array data. As we would not expect heterozygous calls in DH lines all markers with more than 1% heterozygous calls were removed from the panel (34% of all variants). Furthermore, fixated marker positions were also excluded (10% of all variants). Leading to a raw SNP dataset (Figure 1) containing 1,109,642 SNPs (compared to 404,449 variable SNPs with adequate quality (PolyHighResolution [54]) on the high density array [5] (total: 616,201 SNPs)). After the quality filter in the HBimpute step 1,069,959 SNPs remain. Quality control and imputation for the 600k array were performed as described in Pook et al. [39]. As only 1.2% of all markers were imputed this should have a neglectable impact for this study.

### Software

The read-merging and SNP-calling procedure presented in this manuscript are implemented in the R-package HBimpute (available at https://github.com/tpook92/HBimpute). Computing times of the HBimpute pipeline are higher than regular imputation procedures like BEAGLE [31], as the BEAGLE algorithm itself is executed twice and HaploBlocker [39] needs to be applied on the auxiliary imputed SNP dataset (Figure 1). Our pipeline from the raw SNP dataset to the final imputed SNP dataset for chromosome 1 took 107 minutes with 68 minutes spent in BEAGLE for the HB-array pipeline. The HB-seq pipeline took 226 minutes as the haplotype library contained significantly more haplotype blocks that had to be processed in HBimpute. All computing times reported were obtained when using a single core in HBimpute on an Intel(R) Xeon(R) E7-4850 2.00GHz processor. The R-package can be directly be installed within your R session via the following command: **install.packages**(“devtools”) devtools :: **install**_github (“ tpook92/HBimpute”, subdir = “pkg”)

This pipeline is using the software BEAGLE 5.0 as the backend imputation tool (https://faculty.washington.edu/browning/beagle/beagle.html) [31].

## Supporting information

**S1 Table. Predictive ability for the nine maize traits depending on the genotype data used. Details on the individual traits and growing stages (v3-final) can be found in [43].**

**S2 Table. Number of true underlying QTL identified depending on the false discovery rate (FDR).**

## Data Availability Statement

Genomic data for chromosome 1 of the 321 DH-lines that was generated via sequencing with 0.5X read-depth after preprocessing in FreeBayes is available at https://github.com/tpook92/HBimpute. Genomic data for chromosome 1 for the 321 DH-lines that was generated via the 600k Affymetrix® Axiom Maize Genotyping Array is available at https://github.com/tpook92/HaploBlocker. Genomic data for the other chromosomes and raw data are available upon request. All source code underlying the HBimpute step is provided via GitHub (https://github.com/tpook92/HBimpute) and implemented in the associated R-package HBimpute.

## Acknowledgments

The authors thank the German Federal Ministry of Education and Research (BMBF) for the funding of our project (MAZE - “Accessing the genomic and functional diversity of maize to improve quantitative traits”; Funding ID: 031B0882).

