## Supplementary Table S1 & S2 for "Increasing calling accuracy, coverage, and read depth in sequence data by the use of haplotype blocks"

*Table 1 Predictive ability for the nine maize traits depending on the genotype data used. Details on the individual traits and growing stages (v3-final) can be found in Hölker et al. (2019).*

| | HB-seq | HB-seq (overlap) | Array data |
|---|---|---|---|
| Early vigor (v3) | 0.3755 | 0.3804 | 0.3801 |
| Early vigor (v4) | 0.3666 | 0.3732 | 0.3722 |
| Early vigor (v6) | 0.4601 | 0.4639 | 0.4625 |
| Plant height (v4) | 0.5329 | 0.5362 | 0.5347 |
| Plant height (v6) | 0.5559 | 0.5584 | 0.5577 |
| Plant height (final) | 0.7160 | 0.7164 | 0.7160 |
| Days to silking | 0.5416 | 0.5426 | 0.5417 |
| Days to tassel | 0.4835 | 0.4891 | 0.4840 |
| Root lodging | 0.5648 | 0.5695 | 0.5686 |

*Table 2 Number of true underlying QTL identified depending on the false discovery rate (FDR).*

| FDR | Array (600k) | HB-seq | HB-array | BEAGLE | HB-seq (overlap) | HB-seq (large) | Array (50k) | Array (10k) |
|---|---|---|---|---|---|---|---|---|
| 0.40 | 0.21 | 0.24 | 0.28 | --- | 0.18 | 0.22 | 0.13 | --- |
| 0.45 | 0.49 | 0.48 | 0.56 | 0.46 | 0.41 | 0.51 | 0.25 | 0.03 |
| 0.50 | 0.99 | 0.87 | 1.00 | 0.91 | 0.76 | 0.83 | 0.61 | 0.06 |
| 0.55 | 1.51 | 1.46 | 1.62 | 1.53 | 1.35 | 1.40 | 1.11 | 0.26 |
| 0.60 | 2.14 | 2.11 | 2.27 | 2.19 | 1.95 | 2.02 | 1.70 | 0.72 |
| 0.65 | 2.83 | 2.73 | 2.91 | 2.83 | 2.59 | 2.62 | 2.34 | 1.20 |
| 0.70 | 3.55 | 3.44 | 3.62 | 3.57 | 3.29 | 3.29 | 3.06 | 1.72 |
| 0.75 | 4.33 | 4.22 | 4.38 | 4.37 | 4.05 | 4.05 | 3.89 | --- |
